# Epigenetic Profiling of Curcumin on Histone Signatures in Breast Cancer using 3D Network

**DOI:** 10.1101/2024.11.13.623008

**Authors:** Tina Tang, Mikhail Y Berezin, Benjamin A Garcia, Shamim A Mollah

## Abstract

Aberrant epigenetic alterations are implicated in the transformation of normal cells into cancerous ones. Unlike genetic mutations, these dysregulations can be reversible and are potential targets for anticancer drugs. Curcumin, a natural plant-derived compound, has been shown to have anticancer effects and neuroprotective properties via its influence on epigenetic regulation. However, the precise nature of these epigenetic changes and the mechanisms driving them remain largely unexplored. Moreover, there is strong evidence linking the neuroactive properties of chemotherapy drugs in the treatment of breast cancer to chemotherapy-induced peripheral neuropathy (CIPN). Therefore, studying epigenetic regulation of neuroactive compounds in breast cancer is significant in also understanding their mechanisms mediating neuropathy in breast cancer. This study aims to apply a 3D network model to profile the histone codes modified by curcumin as a neuroactive compound and to hypothesize the enzymatic pathways driving these modifications in breast cancer. Using multi-omic data from the NIH LINCS program, we identified two novel histone codes associated with curcumin, mediated through 23 phosphoproteins involved in cell signaling. These signatures were linked to genes expressed in the dorsal root ganglion (DRG), which are characterized in this study. Together, the histone and phosphoprotein profiles represent potential biomarkers for the development of chromatin-targeted therapies in breast cancer, as well as new strategies for managing neuropathy in breast cancer patients.

**GitHub URL** Source code is available at https://github.com/smollahlab/ECHC

## Introduction

Breast cancer is currently the most frequently diagnosed cancer according to the World Health Organization (WHO) with 2.3 million new cases every year. The occurrence and progression of breast cancer is determined by a complex interplay of genetic, epigenetic, and environmental factors. While genetic mutations, such as those in BRCA1 and BRCA2, have been traditionally emphasized in the study of breast cancer etiology, there is now more evidence of the significant role that epigenetic changes play in the cellular transition from a normal state to a cancerous one (1). These epigenetic aberrations can alter gene expression in tumor-related genes and disrupt key pathways such as cell differentiation, survival, migration, and invasion (2). Furthermore, these epigenetic aberrations, unlike genetic mutations, are reversible and represent potential targets for cancer therapy development. However, knowledge about the identity of these epigenetic alterations as well as the mechanisms behind the cause and consequence of them remain largely unknown (3).

The epigenetic markers that this study focuses on are post-translational modifications (PTMs) on histone proteins, one of the structural building blocks that make up chromatin. These PTMs include four different types of covalent modifications on the N-terminal tails of histones: acetylation, methylation, phosphorylation, and ubiquitination (4). Depending on the type, number, and location of these PTMs, which constitute a histone code, they can have different effects on chromatin structure and gene regulation. For example, H3K4me3 promotes the open chromatin state and thus gene transcription; however, H3K27me3 is a repressive mark and indicative of a closed chromatin state (5). These histone PTMs, which are modulated by specialized enzymes termed “readers”, “writers”, and “erasers”, are often the endpoints of protein signaling events following environmental stimuli, such as drug exposure (6). The phosphoproteins involved in these signaling networks can also serve as targets for epigenetic drug therapies (7).

Among the substances studied for their epigenetic activities, curcumin is a natural bioactive compound with pleiotropic biological effects. Derived from the spice turmeric, it has been used in traditional Asian cuisine and medicine for hundreds of years and reported to have an array of pharmacological properties including anti-inflammatory, antioxidant, neuroprotective, and anticancer (8, 9, 10). Curcumin has been shown to suppress breast cancer through the regulation of a multitude of cellular signaling pathways such as cell proliferation, apoptosis, cell-cycle arrest, reactive oxidative stress (ROS), and microRNAs with molecular targets ranging from NF-κB, cyclinD1, P13K, Raf-1, NRF2 and p53 (11, 12, 13, 14, 15, 16, 17, 18, 19). In addition to these mechanisms, curcumin has also been found to influence histone modifications, modulating histone acetylation and methylation in cancer cell lines (5). For example, in one study on triple-negative breast cancer (TNBC), curcumin was shown to inhibit oncogenic EZH2 by reducing H3K27me3 at the promoter for tumor suppressive DLC1, inducing apoptosis in tumor cells (2). However, there is still a need for more studies determining curcumin’s effect on specific histone targets as well as the signaling mechanisms behind its epigenetic regulation.

The neuroprotective effect of curcumin may play an additional, important role in breast cancer treatment as an adjuvant to current chemotherapy treatments. One of the most common side-effects and major limitations in the administration of antineoplastic drugs is chemotherapy-induced peripheral neuropathy (CIPN) (20, 21). Chemotherapeutic drugs, such as oxaliplatin, indiscriminately target healthy cells, including neuronal and glial cells, leading to cell damage and the initiation of neuropathy (20, 21). CIPN symptoms can range from acute, transient sensory abnormalities to chronic pain that persists past cessation of chemotherapy (20). Preliminary evidence links the regulation of neuropathic pain to changes in histone modifications, such as histone acetylation and deacetylation, but the exact effects and mechanisms still require further study (22). Curcumin has been shown to counter neurotoxicity and alleviate neuropathic pain through antioxidative, anti-inflammatory, and epigenetic mechanisms such as the inhibition of histone acetyltransferase (HAT) activity on pronociceptive molecules (23, 24, 25, 26). Several studies on the use of curcumin as an adjunct to oxaliplatin therapy show its effectiveness in protecting against the adverse effects of the platinum-based agent without decreasing its anticancer potential (27, 28, 29). However, more information is required on the epigenetic targets and mechanisms of action behind curcumin’s attenuation of neuropathy.

The goal of this paper is to expand our understanding of curcumin’s influence on histone signatures by using machine-learning techniques to integrate and analyze multi-omics data collected from various cell lines. These signatures will then be used to postulate potential signaling pathways that curcumin utilizes to facilitate its anticancer and neuroprotective effects. This data was compiled as part of the NIH Library of Integrated Network-Based Cellular Signatures (LINCS) program, which aimed to systematically catalog cellular responses to a range of chemical, genetic, and disease perturbagens; curcumin is included within its list of compounds with neuroactive properties (30). Utilizing the 3D data connectivity model developed by Mollah et al. called iPhDNet (31) and it’s generalized version Gen3DNet (32), we aim to profile the histone signatures that encapsulate the curcumin drug response in breast cancer as well as the relevant phospho-signaling proteins that underlie its mechanisms of action. To do this, we will 1) identify the functional modules of histone codes across different cell lines, 2) deduce which phosphoproteins contribute to these histone functional modules in breast cancer, and 3) link these breast cancer signatures to dorsal root ganglion (DRG) genes implicated in neuropathy to hypothesize the potential signaling pathways mediated by curcumin. Using this data-driven approach, we hope to provide deeper insight into curcumin’s epigenetic mechanisms as well as its potential efficacy as a neuroactive compound in breast cancer drug therapies.

## Materials and Methods

### Data Acquisition

Experimental data was downloaded from the LINCS Proteomic Characterization Center for Signaling and Epigenetics (PCCSE) Panorama Repository (https://panoramaweb.org/LINCS). The data was collected from five different cancer cell lines including MCF7 (breast), PC3 (prostate), A375 (skin), A549 (lung), and YAPC (pancreatic), as well as two neural cell lineages: neural progenitor cells (NPC) and astrocytes (33). Each cell line was perturbed by 31 neuroactive drugs, curcumin being one of them, at various concentrations (33). For each drug, there were three biological replicates available, with dimethyl sulfoxide (DMSO) being used as the negative control. Afterwards, the cellular state was captured through two different mass-spectrometry based, proteomic assays that measured phosphoproteins (P100) and global chromatin profiles (GCP) (34, 35). The P100 is targeted against 96 phosphorylated peptides represented in cellular signaling pathways, while the GCP is targeted against 79 combinatorial histone PTMs (33). Both assays were each acquired at Level 3, which is preprocessed to be log 2 transformed and normalized. The P100 was collected at 3 hours after treatment while the GCP was collected at 24 hours after treatment.

In addition to the proteomic assays, we also acquired L1000 RNA transcriptomic data for the MCF7 (breast cancer) cell line treated with curcumin. The L1000 assay is a targeted gene expression assay against 978 landmark genes that are representative of the entire transcriptome (36). These were obtained at Level 5, which is preprocessed to be a differential expression signature called a CD-coefficient using the characteristic direction method developed by Ma’ayan et al (37). The L1000 assays were downloaded at 6 hours and 24 hours after treatment with curcumin from the SigCom LINCS web server (https://maayanlab.cloud/sigcom-lincs). A complete list of all data files used is found in **Supplementary Table 1**.

### Data Preprocessing

For each of the 31 drugs in the proteomic assays (P100 and GCP), the three replicates were averaged and used to impute any missing values. The differential fold change was then calculated by taking the fold change of each perturbed phosphoprotein or histone with respect to the control (DMSO). The final preprocessing result is two data matrices: a histone matrix [31 neuroactive drugs x 52 histone marks] and a phosphoprotein matrix [31 neuroactive drugs x 72 phosphopeptides]. For the identification of histone modules across different cell lines, another histone matrix was created that consisted of histone marks perturbed by curcumin in each cell line [7 cell lines x 52 histone marks].

### Gen3DNet Data Analysis Pipeline

Generic 3D Network (Gen3DNet) (32), first derived from Integrated Phosphoprotein Histone Drug Network (iPhDNet) (31), is the data analysis pipeline used in this study that was developed by Mollah et al. Gen3DNet (32) is the generalized version of iPhDNet (31) that consists of two parts: a non-negative matrix factorization (NMF) and a partial least square regression (PLSR) component. It takes two data inputs: a histone matrix and phosphoprotein matrix. **Figure 1** summarizes this workflow below.

**Figure 1:**
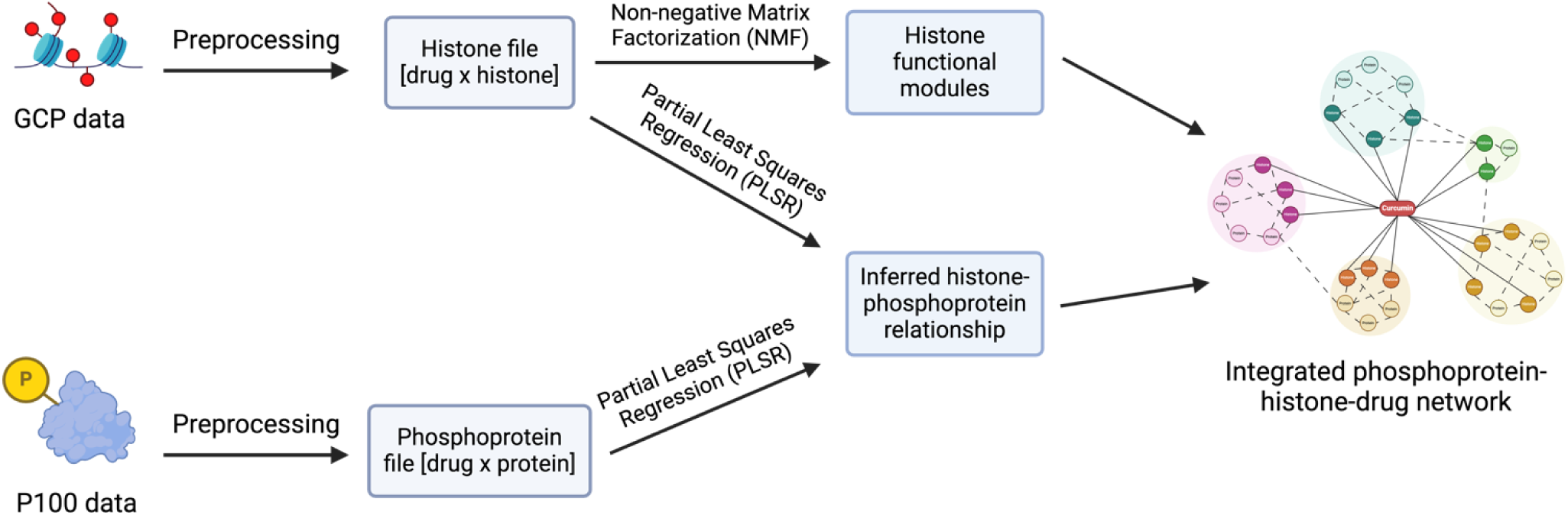
Gen3DNet Workflow. This workflow shows how an integrated three-dimensional network connecting drugs, phosphoproteins, and histone marks can be generated from two proteomic assays, GCP and P100, using Gen3DNet.

In the first component, NMF is used to identify the histone signatures (31). The goal of NMF is to extract a smaller number of latent factors (**k**) that can explain the data; these latent factors can be used to cluster the data into **k** modules. It does this by factoring the original matrix **A** (the histone matrix; N x M) into two matrices, **W** (basis matrix; N x k) and **H** (loading matrix; k x M), where the product of **W** and **H** approximate **A** (**A ≈ WH,** where **W, H ≥ 0**). The coefficients of the basis matrix signify the association of each histone code with a cluster, while the loading or mixture coefficients represent the connection of each drug (or cell line) to a cluster. This ultimately can be used to group a set of drugs with a collection of histone codes, which we refer to as a histone signature, to characterize the drug response.

In the second component, PLSR is used to create the histone-peptide interaction network (31). PLSR is a multivariate, linear regression model that finds the fundamental relations, known as the latent variables, between two matrices: **X** (predictor matrix) and **Y** (response matrix). In our application, the histone matrix is modeled as the response matrix (**Y**) and the phosphoproteins are the predictor variables (**X**). PLSR is useful in cases where there are many predictor variables with high collinearity, such as in this case where the phosphoproteins are highly correlated with each other. The resulting regression coefficients thus represent the contribution of each phosphoprotein to a specific histone code and the strength of their interaction. Finally, a t-test (**p < 0.01**) was used on these PLSR coefficients to determine the statistically significant phosphoproteins.

### Integrated Histone-Phosphoprotein-Drug Network

To generate the three-dimensional network of interactions, the NMF histone-drug signatures for curcumin were integrated with the predicted PLSR histone-phosphoprotein relationships. This was visualized using Cytoscape (38) where the histones, phosphoproteins, and drugs made up the individual nodes, and the NMF or PLSR coefficients were the edges connecting histones to curcumin or histones to phosphoproteins, respectively.

### Experimental Design and Statistical Rationale

All experimental assays were previously generated as part of the NIH LINCS program (33, 34). Seven different cell lines were perturbed by 31 neuroactive drugs with three biological replicates taken for each drug; after drug perturbation, changes in phospho-signaling and histone modifications were measured via the P100 and GCP assays, respectively (33, 34, 35). Utilizing the R software package Gen3DNet (32), we identified drug-histone signatures and predicted histone-phosphoprotein interactions to construct a 3D connectivity map among curcumin, phosphoproteins, and histone codes. We excluded any samples that had no recorded measurements to create a uniform list of proteins and histone marks across cell lines. In this study we had two independent groups: a treatment group and control group; randomization and blinding were not relevant to our study. After 3D network analysis, we applied a t-test (p-value < 0.01) to retain the statistically significant 39 phosphoproteins from P100 and 2 histones from GCP.

## Results

### Two functional histone modules identified across different cancer and neural cell lines

To better understand the context of curcumin’s cellular response, we utilized NMF to identify two functional modules in relation to the five cancer and two neural cell lines. Our results are displayed as heatmaps of basis and mixture components from NMF that correspond to each histone code and cell line’s membership to a histone signature, respectively (**Figure 2A**). These two histone modules characterize the cellular response state to curcumin. Thirty unique histone codes are associated with the first module and nineteen with the second module. The first module is made up of the cell lines MCF7 (breast cancer), PC3 (prostate cancer), NPC (neural progenitor cell), and astrocyte; while the second module consists of YAPC (pancreatic cancer), A375 (skin cancer), and A549 (lung cancer). Seeing how both MCF7 and PC3 are clustered together and are cancers affected by hormone dysregulation gave us good intuition that there were similarities in how curcumin affects the epigenetic landscape in both diseases. Likewise, since MCF7 was also grouped together with NPC and astrocyte, this indicates that curcumin may have related epigenetic mechanisms in breast cancer and neurological pathologies. This is further supported by the fact that MCF7 shares ectodermal origin with NPC and astrocyte cells (39, 40, 41). The detailed summary of histone modification and cell line clustering results can be found in **Supplementary Table 2**.

**Figure 2:**
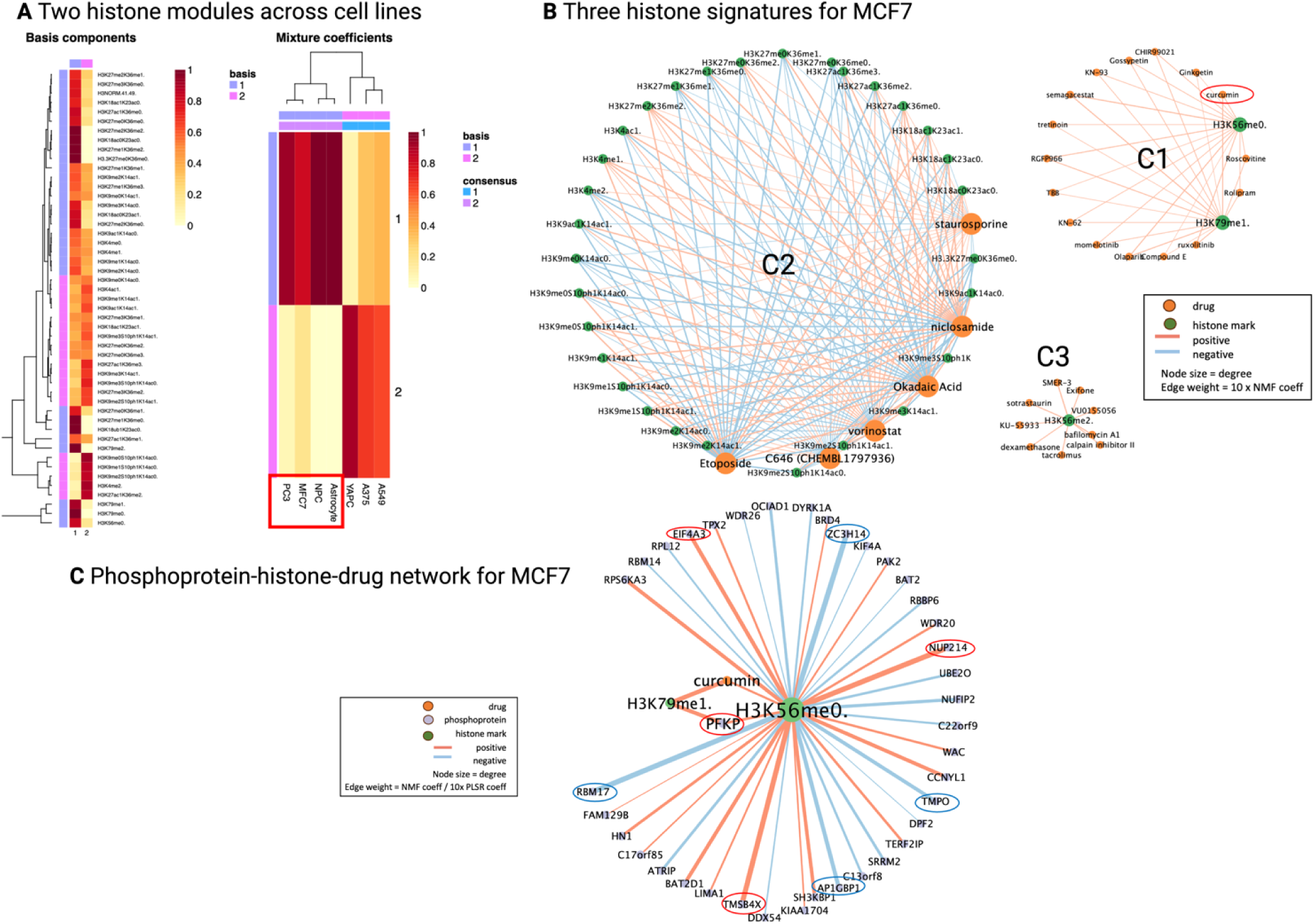
Network analysis results for predicted histone-phosphoprotein interactions associated with curcumin. (A) Two histone modules characterized across seven cell lines. The basis components signify the strength of each histone code’s membership to a functional module, while the mixture coefficients reveal the contribution of each cell line towards a histone signature. (B) Three histone signatures categorized for the MCF7 cell line. While the orange and green nodes represent neuroactive drugs or histone marks, respectively, the light red and blue edges represent elevated or reduced histone response levels. (C) Histone-phosphoprotein-drug network for the MCF7 cell line perturbed by curcumin. Curcumin is signified by the orange node, while phosphoproteins and histone marks are shown in purple and green, respectively. Elevated and reduced phosphoprotein/histone levels are represented as light red and blue edges. The top enriched phosphoproteins in this network are circled in red and blue standing for the top elevated and reduced peptides, respectively.

### Phosphoprotein-histone associations mediated by curcumin in breast cancer

To further decipher curcumin’s effect in breast cancer, we used network analysis to identify specific histone codes perturbed by curcumin and construct interactions between those histone codes and signaling phosphoproteins. First, NMF was used on the MCF7 GCP data to cluster the histone codes and neuroactive drugs into three distinct signatures, with curcumin being found in the second cluster along with two histone codes: H3K79me1 and H3K56me0 (**Figure 2B**). **Supplementary Table 3** recaps each histone signature with their respective drug groupings and associated drug pathways.

Next, PLSR was utilized on the P100 phosphoproteins and GCP histones to predict phosphoprotein-histone relations that are then integrated with the histone signatures mentioned previously. PLSR model performance is shown in **Supplementary Figure 1**. After filtering for the statistically significant phosphoprotein interactions which resulted in 40 unique associations between 39 distinct phosphoproteins and the two histone codes, the outcomes are visualized in a three-dimensional network that connects phosphoproteins to histones to curcumin (**Figure 2C**). All 40 predicted curcumin histone-phosphoprotein relations are detailed in **Supplementary Table 4**. The top enriched phosphoproteins in this network, which are circled on the Cytoscape-generated network, include: ZC3H14, NUP214, RBM17, TMSB4X, AP1GBP1, EIF4A3, PFKP, and TMPO.

### Two histone modifications connected to curcumin in breast cancer

A literature search on H3K56me0 and H3K79me1 reveals the histone PTMs from the Gen3DNet analysis found to be significantly connected to curcumin. The results are summarized in **Supplementary Table 5**. Both histone marks were found to be elevated in our study; to our best knowledge this is the first study of these histone codes in relation with curcumin. The first histone mark, H3K56me0, is a novel mark with limited data on its biological function. However, past studies have characterized H3K56me1 as an important facilitator in DNA replication by serving as an important docking site for nuclear machinery (42), suggesting that the absence of methylation of K56 in H3K56me0 may lead to cell cycle disruption. This would help support the possible case of curcumin acting to induce cell cycle arrest in breast cancer cells to suppress tumor growth.

The second histone mark, H3K79me1, is associated with transcriptional activation along with its counterparts, H3K79me2/3 (43, 44). One study reports that the transition from mono-methylation to di- and tri-methylation at H3K79 is correlated with low- to high-level gene transcription activity (45). While there are no studies directly connecting H3K79me1 to breast cancer, there is evidence that H3K79me2 is positively correlated with breast cancer gene expression (46). Additionally, inhibition of H3K79 methylation by targeting DOT1L, the only known H3K79 methyltransferase, suppresses breast cancer proliferation (47, 48, 49). Though H3K79me1 is elevated in our study, it is possible that the role of mono-methylation in cellular regulation, and/or its location in the genome, is distinct from that of H3K79me2 and may still contribute to breast cancer repression.

In addition to their role in breast cancer, H3K79 methylation and DOT1L activity have also been found to be important for neurological function. For example, H3K79 methylation is necessary for neuronal development and regulation (50). DOT1L has also been discovered to help maintain proper cortical and cerebellar development (51). These findings suggest that H3K79 methylation may be a possible epigenetic target in the cellular mechanisms by which curcumin acts as a neuroactive drug.

### Functional profiling of enriched phosphoproteins associated with histones PTMs and comparison to neural gene lists

To augment our findings on the enriched P100 phosphoproteins obtained from our network analysis, we incorporated the L1000 gene expression data from the MCF7 cell line perturbed by curcumin. Of the 39 unique phosphoproteins identified as significant by PLSR, 23 of those matched to genes captured in the L1000 assay (**Figure 3A**). Given that the phosphoproteins were measured at 3 hours and RNA expression at 6 and 24 hours, we decided to compare the direction (positive or negative) of the L1000 CD-coefficient with the direction of the P100 fold change. For both CD-coefficient and fold change, a positive value signifies an increase in expression whereas a negative value indicates a decrease in expression. As expected, there were more L1000 genes sharing the same direction with the P100 phosphoproteins at 6 hours than at 24 hours, with 15 of the 23 genes matching at 6 hours but only 6 genes matching at 24 hours (four of these genes matched at every time point) (**Figure 3B**). Thereafter, we narrowed down our list of enriched phosphoproteins to these 23 that had corresponding gene expression values.

**Figure 3:**
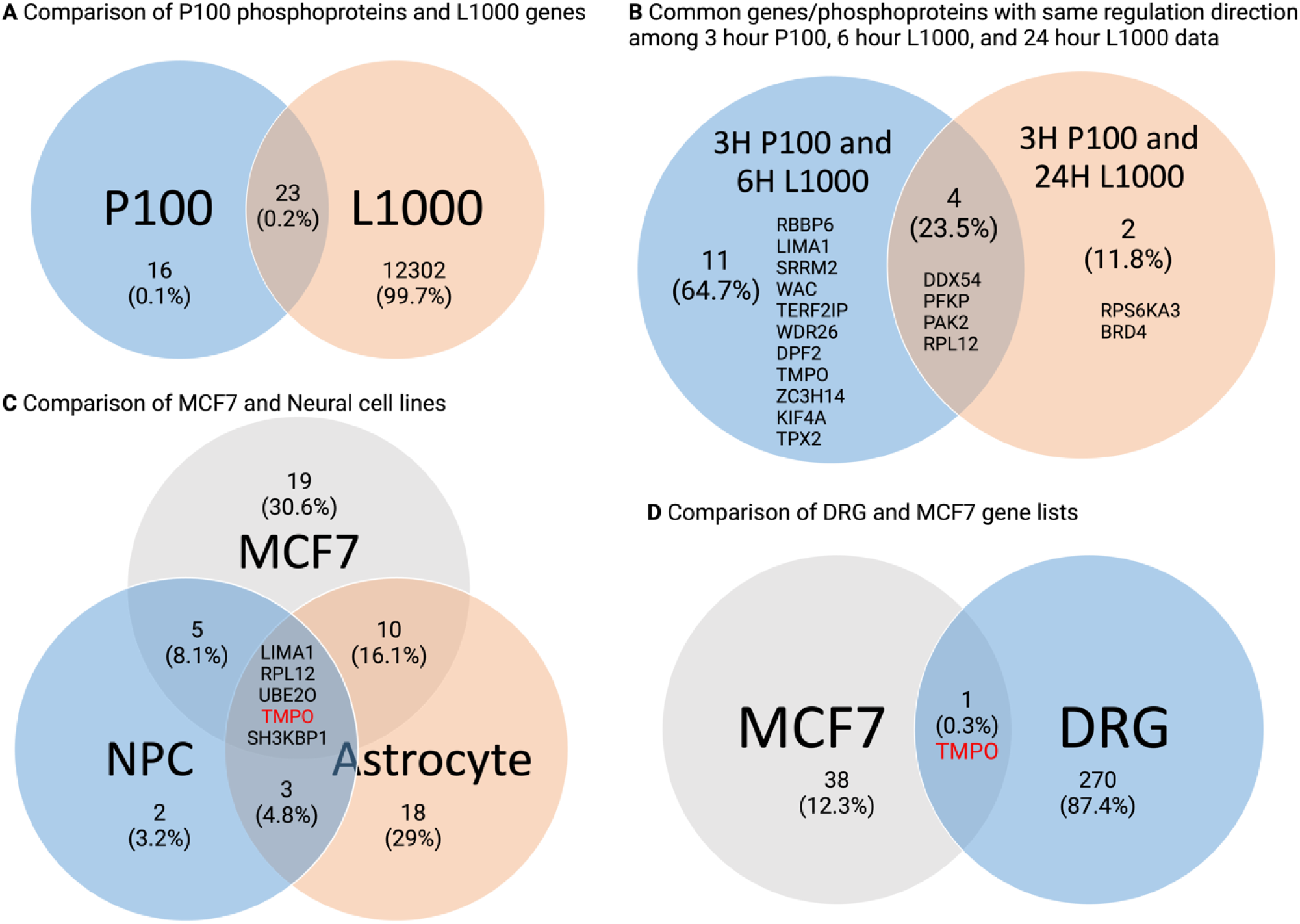
P100 (phosphoprotein), L1000 (transcriptomic), and neural cell gene list (NPC, Astrocyte, DRG) comparisons. Enriched P100 phosphoproteins of interest were compared (A) to the L1000 genes list, (B) across different time points by regulation direction (with 11 genes matching directions at 3 and 6 hours, 2 genes matching at 3 and 24 hours, and 4 genes matching across all time points), (C) to the phosphoproteins associated with curcumin in neural cell lines (NPC and astrocyte), and (D) to the list of significant DRG neuropathic genes.

To determine which of the 23 phosphoproteins might be meaningful targets for drug interventions, we consulted several different resources and analysis tools. This included a literature search for previous studies connecting to breast cancer, functional enrichment analysis, gene annotation databases such as GeneCards (52), and additional neurological gene lists. Previous literature connects the expression of several of these enriched phosphoproteins to cancer. For example, RBM17, an RNA binding motif protein, is overexpressed in many cancer types and associated with drug resistance and shortened survival (53, 54). Similarly, TMPO is also upregulated in multiple cancer cell lines, including breast cancer (55). Both proteins are reduced in the P100 data, corresponding to curcumin’s potential anticancer activity. On the other hand, high expression of RPS6KA3 aka RSK2, a serine/threonine kinase, showed more favorable survival in breast cancer patients (56) and is elevated in our study. The functional significance and results from the LINCS data of the top enriched phosphoproteins are further profiled in **Supplementary Table 6**.

Overall, the most enriched biological processes identified by Metascape for these 23 genes included regulation of cytoskeleton organization, axon guidance, RNA splicing, and cell cycle (**Figure 4A**). Several of the top enriched phosphoproteins are associated with these biological functions, including EIF4A3 and RPS6KA3 in nervous system development, EIF4A3 and RBM17 in mRNA splicing, NUP214 and TMPO in cell cycle regulation (61). An extensive and detailed list of the top enriched processes and their correlated genes is provided in **Supplementary Table 7**.

**Figure 4:**
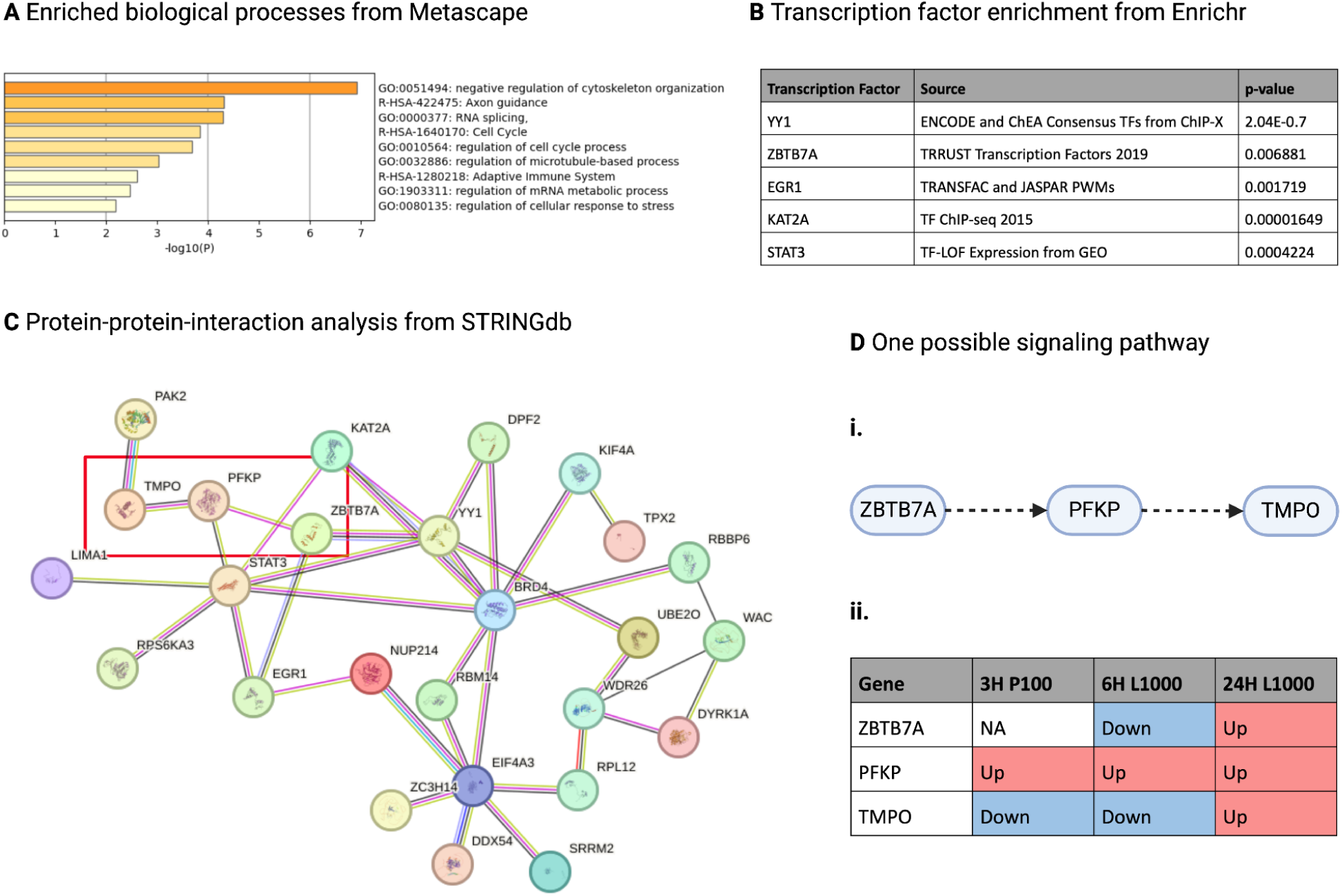
Gene list enrichment analysis results. (A) Metascape pathway enrichment analysis. (B) Enrichr transcription factor enrichment analysis. (C) STRINGdb protein-protein-interaction analysis. (D) Example of potential drug signaling pathway analyzed across time points.

Since curcumin is a known pleiotropic drug with neuroactive properties, we further investigated which of these phosphoproteins of interest overlap with genes that have been implicated in neural cell lines and neuropathy. While DRG neurons are the primary target of chemotherapy-induced neurotoxicity, astrocyte activation has been also shown to play a role in the initiation of CIPN, leading to our interest in studying a range of different types of neural cells (20, 21, 27, 57, 58, 59). To do this, we first observed to see if shared phosphoproteins existed between the top enriched proteins associated with curcumin from MCF7 and those identified in NPC and astrocyte cell lines (**Figure 3C**). There were five genes that coincided with all three cell lines: LIMA1, RPL12, UBE20, SH3KBP1, and TMPO. We also cross-referenced with a list of genes obtained from the DRG of mice that had been treated with oxaliplatin, a chemotherapeutic agent used in cancer treatment, that had been implicated in the pathophysiology of CIPN (https://www.ncbi.nlm.nih.gov/geo/query/acc.cgi?acc=GSE286387) (60). Of the 271 differentially expressed significant DRG genes (FDR<0.05, fold-change > 1.5), the TMPO phosphoprotein was found to be enriched (**Figure 3D**).

### Protein-protein interaction and transcription factor analysis exhibited potential signaling pathways

To identify possible gene regulatory networks, we inferred signaling pathways for curcumin by exploring connections between the phosphoproteins-of-interest and transcription factors (TFs). We used Enrichr, a transcription factor enrichment tool, to identify associated transcription factors from the gene list of 23 enriched genes (62). Based on probability scores and biological significance in breast cancer, we selected five transcription factors: YY1, ZBTB7A, EGR1, KAT2A, and STAT3 (**Figure 4B**). We then used the STRING database, a protein-protein interaction (PPI) tool (63), to predict interactions among those transcription factors and the 23 enriched phosphoproteins which function as possible TF targets. The resulting protein-protein interaction network is displayed in **Figure 4C**.

When proposing potential signaling pathways, we focused on relationships linking TFs to the top enriched phosphoproteins from the Gen3DNet network analysis. One example of a possible signaling pathway, involving the transcription factor ZBTB7A influencing the phosphoproteins PFKP and TMPO downstream, is outlined in **Figure 4C** and depicted in **Figure 4Di**. We observed each gene in this pathway across three time points to determine if the expression data, either increased (“up”) or decreased (“down”), can match the role that each gene plays in the pathway and in breast cancer to determine the pathway’s feasibility in breast cancer suppression. The expression of PFKP is increased at all available time points after curcumin exposure, while the expression of TMPO and ZBTB7A are initially decreased (note there is no P100 data available for ZBTB7A since it is not a phosphoprotein) (**Figure 4Dii**). However, both ZBTB7A and TMPO switched from being down-regulated to being up-regulated at 24 hours, implying that perhaps the influence of curcumin declines within 24 hours after first exposure

We characterized additional potential pathways from the PPI analysis in **Supplementary Table 8**. These include a similar signaling pathway involving transcription factor STAT3 with PFKP and TMPO; however, since the upregulation of STAT3 is known to promote breast cancer malignancy (64), its elevation in our study makes its anti-oncogenic potential with curcumin more ambiguous. Another possible pathway linking transcription factor EGR1, nucleoporin NUP124, and RNA helicase EIF4A3 was also considered. EGR1 is upregulated in the L1000 data, corresponding to breast cancer suppression (65). EIF4A3 was also elevated in our study; however, it is the inhibition of EIF4A3 that has been shown to suppress breast cancer proliferation and metastasis (66, 67, 68). Given these evaluations and our interest in the TMPO phosphoprotein, the signaling pathway of ZBTB7A to TMPO was selected as the most pertinent and probable mechanism of action for curcumin in breast cancer.

## Discussion

This study serves as an example of how the application of a 3D network tool (Gen3DNet) (32) on complex, multi-omic data allows for the extraction of a smaller number of meaningful features that can be further examined and constructed into data-driven hypotheses. This allows us to theorize potential epigenetic signaling pathways for curcumin in breast cancer. In this study, we primarily analyzed proteomic data from the MCF7 cell line which resembles the Luminal A (ER+, PR+) breast cancer subtype. The findings from this study have potential applications in identifying histone PTM and phosphoprotein changes corresponding to curcumin treatment in this specific subtype of breast cancer.

From our profiling of curcumin’s epigenetic mediation, thymopoietin (TMPO) emerged as a phosphoprotein of interest. As one of the top enriched phosphoproteins in our histone-phosphoprotein network analysis, TMPO is a structural protein found in the nucleus and involved in several cellular processes including gene transcription, DNA replication, and cell cycle control (69). It has been implicated in cancer progression and found to be upregulated in several cancers, including breast cancer; its depletion is associated with cell cycle arrest and apoptosis in cancer cell lines (55). In our study, TMPO is downregulated, potentially corresponding to curcumin’s anti-cancer activity. Furthermore, as neuropathy is a common side-effect in chemotherapy treatments for breast cancer, it is important to consider curcumin’s dual role in the mediation of CIPN in addition to breast cancer suppression. Based on the results of this study, TMPO may play a key role in both. When drawing comparisons with the neural cell lines, TMPO was among the shared enriched phosphoproteins between the breast cancer (MCF7), neural progenitor cell (NPC), and astrocyte cell lines. Moreover, TMPO was the sole gene that matched with the DRG genes linked to CIPN (60). Given its neurological importance, as well as its involvement in the cell cycle and cancer, we present TMPO as a possible key player in curcumin’s epigenetic mechanism of action.

Among the possible pathways that involve TMPO, the most viable pathway involves the transcription factor ZBTB7A and phosphoprotein PFKP. ZBTB7A is a zinc finger protein that functions as a transcriptional repressor for a wide range of genes involved in cell proliferation and differentiation, including PFKP; it has been reported to accelerate breast cancer progression (70, 71). In our study, ZBTB7A is downregulated in the L1000 expression data captured at 6 hours after curcumin exposure which correlates with cancer suppression. PFKP, a key enzyme in glycolysis regulation, can play a role in metabolic reprogramming in some cancers, including bladder, breast, and lung; high levels of PFKP are associated with poor prognosis in cancer patients (72, 73). PFKP is upregulated in our study which correlates with ZBTB7A inhibition but does not, however, correspond with breast cancer repression according to the current literature. There may be compensatory effects from other regulators recruited by PFKP that may account for this. There is also no well-documented relationship between PFKP and TMPO at the present, though both proteins can be localized to the nucleus; further studies are needed to validate this connection (74). In addition, to this date there are no studies on the relationships between PFKP and TMPO with the histone mark H3K56me0 (and similarly between PFKP and H3K79me1).

Furthermore, this is the first study connecting these histone signatures to curcumin treatment in breast cancer. These signatures include the function and regulation of the histone PTMs identified as part of curcumin’s histone-drug signature, the predicted phosphoprotein-histone relationships, as well as the signaling pathways for gene regulation connecting breast cancer and neuropathy. To be able to effectively evaluate and validate all these new inferred relationships, more sequence-driven assays, such as ATAC-seq (75), ChIP-seq (76), and CUT&RUN (77), would be required. This would allow us to identify the location of histone peaks for the histone PTMs of interest as well as perform sequence-driven TF-motif analysis. The next step in determining causality would be experimental validation in a biological model, such as performing a knockout of TMPO and measuring breast cancer progression.

This study also brought to light some limitations that arise when answering biological questions regarding causal interactions between genetic and epigenetic factors. The biggest limitation encountered during the analysis process was the lack of consistent time points across assays. The difference in collection time between the P100 and GCP assays left some ambiguity in the predicted phosphoprotein-histone relationships, as histone PTMs are very transient and can change quickly between 3 and 24 hours. We attempted to provide additional data points with the L1000 assays at the time points of 6 and 24 hours, though the correlation between RNA expression and phosphoprotein level may also be limited. Ideally, each type of assay would be collected at the same time point, across multiple time points. Without this data consistency, identifying a mechanism of action across multi-omic components and providing evidence for causation is more challenging.

In conclusion, in this study we provided a systematic approach to profile histone signatures and generate data-driven hypotheses on epigenetic signaling mechanisms induced by curcumin. We achieved this by using the network analysis pipeline of Gen3DNet to identify drug-histone signatures and predict histone-phosphoprotein interactions, with the final outcome being a connectivity map among curcumin, phosphoproteins, and histone codes. To our knowledge this is the first study to analyze and profile upwards of 50 histone codes in relation to curcumin in breast cancer and link novel histone PTMs with curcumin treatment. In this study we also found possible connections between curcumin’s epigenetic targets in breast cancer and neurological disorders, suggesting potential common epigenetic pathways utilized by these diseases. Overall, our study elucidated possible new therapeutic targets for curcumin as an epigenetic modulator for breast cancer and other diseases.

## Supporting information

Supplemental files for tables

## Acknowledgements

This work was supported by Washington University in St. Louis School of Medicine Genetics departmental funding and NIH/NHGRI U01HG013227-01. DRG gene list generation was supported by NIH/NCI R01CA208623, R21CA269099. We want to thank the NIH LINCS consortium for making these data available.

## Author Contributions

**Tina Tang:** Software implementation, Investigation, Data curation, Writing- Original draft preparation, Visualization. **Mikhail Berezin:** Resources- DRG gene list, Writing- Review. **Benjamin Garcia:** Writing- Review. **Shamim Mollah:** Conceptualization, Methodology, Software development, Writing- Review & Editing, Supervision, Funding Acquisition.

## Data Availability

Code and analysis results are available on GitHub: https://github.com/smollahlab/ECHC. The Gen3DNet R package is also available on Github: https://github.com/smollahlab/gen3DNet.

